# Developmental loci harbor clusters of accelerated regions that evolved independently in ape lineages

**DOI:** 10.1101/226019

**Authors:** Dennis Kostka, Alisha K. Holloway, Katherine S. Pollard

## Abstract

Some of the fastest evolving regions of the human genome are conserved non-coding elements with many human-specific DNA substitutions. These Human Accelerated Regions (HARs) are enriched nearby regulatory genes, and several HARs function as developmental enhancers. To investigate if this evolutionary signature is unique to humans, we quantified evidence of accelerated substitutions in conserved genomic elements across multiple lineages and applied this approach simultaneously to the genomes of five apes: human, chimpanzee, gorilla, orangutan, and gibbon. We find roughly similar numbers and genomic distributions of lineage-specific accelerated regions (linARs) in all five apes. In particular, apes share an enrichment of linARs in regulatory DNA nearby genes involved in development, especially transcription factors and other regulators. Many developmental loci harbor clusters of nonoverlapping linARs from multiple apes, suggesting that accelerated evolution in each species affected distinct regulatory elements that control a shared set of developmental pathways. Our statistical tests distinguish between GC-biased and unbiased accelerated substitution rates, allowing us to quantify the roles of different evolutionary forces in creating linARs. We find evidence of GC-biased gene conversion in each ape, but unbiased acceleration consistent with positive selection or loss of constraint is more common in all five lineages. It therefore appears that similar evolutionary processes created independent accelerated regions in the genomes of different apes, and that these lineage-specific changes to conserved non-coding sequences may have differentially altered expression of a core set of developmental genes across ape evolution.

## Introduction

Accelerated sequence evolution is a hallmark of both positive selection and loss of constraint. Therefore, the comparative genomic signature of sequence change in a lineage of interest compared to conservation in other lineages has been used to identify genome sequences that are candidates for explaining evolution of lineage-specific traits. The requirement of conservation in other lineages serves two purposes. First, it suggests functional constraint, which enables genome-wide scans to focus on regions where accelerated evolution is most likely to have meaningful consequences. This is particularly helpful for discovering accelerated non-coding elements without clear-cut functional annotations, and less so for tests of accelerated evolution in well-annotated genes (Kosiol, Vinar et al. 2008). The second reason to focus on regions that are otherwise conserved is the higher power to detect a shift in evolutionary rate, including shifts from conserved to neutrally evolving (e.g., loss of function) or weak positive selection (Pollard, Salama et al. 2006).

This approach was applied genome-wide to identify Human Accelerated Regions (HARs) that experienced significantly more substitutions than expected in the human lineage since divergence from the common ancestor with chimpanzees (Pollard, Salama et al. 2006, Prabhakar, Noonan et al. 2006, Bird, Stranger et al. 2007, Bush and Lahn 2008). It has also been applied to other lineages, including fruit flies (Holloway, Begun et al. 2008), the common ancestor of therian mammals (Holloway, Bruneau et al. 2016) and the common ancestor of bats (Booker, Friedrich et al. 2016). The ~2,700 HARs identified to date are mostly non-coding, and many have epigenomic signatures suggestive of enhancer function in human cells. To date, 62 out of 92 HARs that were prioritized based on evidence of a regulatory role (67.4%) showed enhancer activity in transient transgenic reporter assays in mouse or fish embryos, and seven HARs are known to drive different expression patterns with the human compared to orthologous chimpanzee sequence (Hubisz and Pollard 2014). Consistent with this role in developmental gene expression, HARs are enriched in genomic loci with transcription factors and other genes involved in regulation of development (Capra, Erwin et al. 2013, Kamm, Pisciottano et al. 2013, Gittelman, Hun et al. 2015). HARs also occur at a higher rate at the distal ends of chromosomes, where sequence divergence is generally higher (Pollard, Salama et al. 2006).

We were curious if the prevalence of lineage-specific accelerated regions (linARs) and their association with developmental gene regulation is unique to humans, or if chimpanzees and other primates share this genomic feature. Previous work identified linARs that are accelerated across primates as a clade (i.e., with substitutions in multiple primate lineages), and these were indeed mostly non-coding and associated with developmental loci (Lindblad-Toh, Garber et al. 2011). With more genomes now available, it is possible to query each primate lineage individually for linARs and to compare patterns across species. To do so requires a statistically rigorous method for assessing evidence of acceleration in each lineage for each of many genomic regions. We solved this problem by implementing a model selection procedure that uses likelihood ratio statistics to evaluate support in the multiple sequence alignment of a given genomic region for a partially nested set of models with combinations of acceleration or no acceleration in different lineages.

In addition to testing for acceleration, our model selection procedure also enables evaluation of evidence for GC-biased gene conversion (gBGC) in different lineages. gBGC is a recombination associated process that mimics positive selection by increasing the rate of fixation of GC alleles, while decreasing the rate of fixation of AT alleles. Previous analyses of HARs found evidence of gBGC, especially in the fastest evolving HARs and in regions of high recombination in modern humans (Pollard, Salama et al. 2006, Galtier and Duret 2007, Lindblad-Toh, Garber et al. 2011, Kostka, Hubisz et al. 2012). Using the weak-mutation model of Kostka et al. (Kostka, Hubisz et al. 2012), our method therefore evaluates each conserved genomic region with a collection of models that includes all combinations of unbiased acceleration (loss of constraint or positive selection), GC-biased acceleration (gBGC), or no acceleration in each lineage.

We applied this approach to whole-genome alignments and identified 5,916 conserved elements with evidence of accelerated substitutions in at least one of five apes: human (*Homo sapiens*), chimpanzee (*Pan troglodytes*), gorilla (*Gorilla gorilla gorilla*), orangutan (*Pongo pygmaeus abeii*), and gibbon (*Nomascus leucogenys*). These ape linARs are roughly equal in number across species (taking differences in data quality of genome assemblies into account), mostly non-coding, and enriched nearby genes involved in regulation of development. Interestingly, a number of developmental loci harbor distinct linARs that are accelerated in different apes, suggesting that shared developmental pathways have experienced independent bursts of regulatory evolution across distinct ape lineages. Unbiased acceleration, consistent with positive selection or loss of strong constraint, is more prevalent than gBGC in all species, although each ape has a substantial minority of linARs (approximately 26%) that are GC-biased. These findings clarify that linARs are not a human-specific phenomenon.

## New Approaches

### Testing for lineage-specific acceleration across multiple lineages

We developed a statistical procedure to test an alignment *X* for evidence of accelerated substitution rate in any combination of lineages in a phylogenetic tree. Let *L*_0_(*ϑ*_0_, *X*) be the log-likelihood of alignment *X* under a null model of DNA evolution where the parameters *ϑ*_0_ encode the phylogenetic model (tree topology, branch lengths, and DNA substitution rate matrix) without acceleration in any lineage. In practice, *ϑ*_0_ will typically be estimated by scaling a genome-wide tree model to have the maximum likelihood substitution rate and GC-content of *X* (ignoring the species for which acceleration will be tested); that is, it will be estimated without changing tree topology or otherwise altering relative rates of different types of substitutions. When applied to conserved elements, this rescaling means that the null model is conservation in all lineages, including the lineage of interest. We compare this null model with no lineage-specific acceleration to a partially nested series of alternative models ***ϑ***_*A*_ with log-likelihoods *L_Ai_*(*ϑ_Ai_, X*), where the set ***ϑ***_*A*_ = {*ϑ_Ai_*} has one parameter combination *ϑ_Ai_* for each of the possible ways to have of unbiased and/or GC-biased acceleration in the lineages of interest. GC-biased acceleration allows us to detect elements that are accelerated due to biased gene conversion and mismatch repair (Pollard, Salama et al. 2006, Galtier and Duret 2007, Lindblad-Toh, Garber et al. 2011, Kostka, Hubisz et al. 2012), whereas unbiased acceleration capture loss of constraint and positive selection. For example, a specific *ϑ_Ai_* could code for GC-biased acceleration on the human branch and unbiased acceleration on the branch leading to gibbon. Our goal is to find the best alternative model and test if it fits the multiple sequences alignment for a genomic region significantly better than the null model.

### An efficient model selection algorithm

One approach to find the best alternative model (with parameters *ϑ_Ai_* representing acceleration on at least one lineage) is to enumerate all alternative models, compute the likelihood of each one, and select the one with the most significant improvement in likelihood over the null model (with parameters *ϑ*_0_). This can be achieved using likelihood ratio tests (LRTs), penalized based on the number of parameters (lineages with acceleration) to avoid over-fitting. In applications with a large number of alternative models and alignments, it is computationally prohibitive to compare all possible models.

We therefore designed a forward model selection algorithm that takes advantage of the hierarchical structure of the set of alternative models (**Figure 1a–b**). The alternative models are partially nested; for any *i* (except the full model with biased and unbiased acceleration on all lineages) there is at least one model with parameters *ϑ_Aj_* that contains the model with parameters *ϑ_Ai_* as a special case. For example, the model with unbiased acceleration on the human branch is nested within the model with unbiased acceleration on both the human and chimpanzee branches, because our parameterizations for acceleration include no acceleration as a special case (see below). Instead of scoring all models, we start by calculating log-likelihood ratio test statistics *T_i_*(*X*) = *2*(*L_Ai_*(*ϑ_Ai_, X*) – *L*_0_(*ϑ*_0_, *X*)) for the simplest alternative models (acceleration on one branch, either biased or not), and then we proceed to more complex models *ϑ_Aj_* including one extra parameter only if *T_j_*(*X*)–*T_i_*(*X*) =: *Δ_ij_*(*X*) > *c* for at least one *i*; we note that following this approach the model with parameters *ϑ_Ai_* is nested in the model with parameters *ϑ_Aj_*. The constant *c* is based on the asymptotic distribution of *Δ_ij_*(*X*) under *ϑ*_0_ and chosen such that *Pr*(*Δ_ij_*(*X*) > *c* | *ϑ*_0_) = *p*, where *p* is a small number (we use *p* = 0.01 in this study). Thus, the more complex model with paramters *ϑ_Aj_* is only selected if there is evidence that it fits the data significantly better than the simpler model with paramters *ϑ_Ai_*.

**Figure 1.**
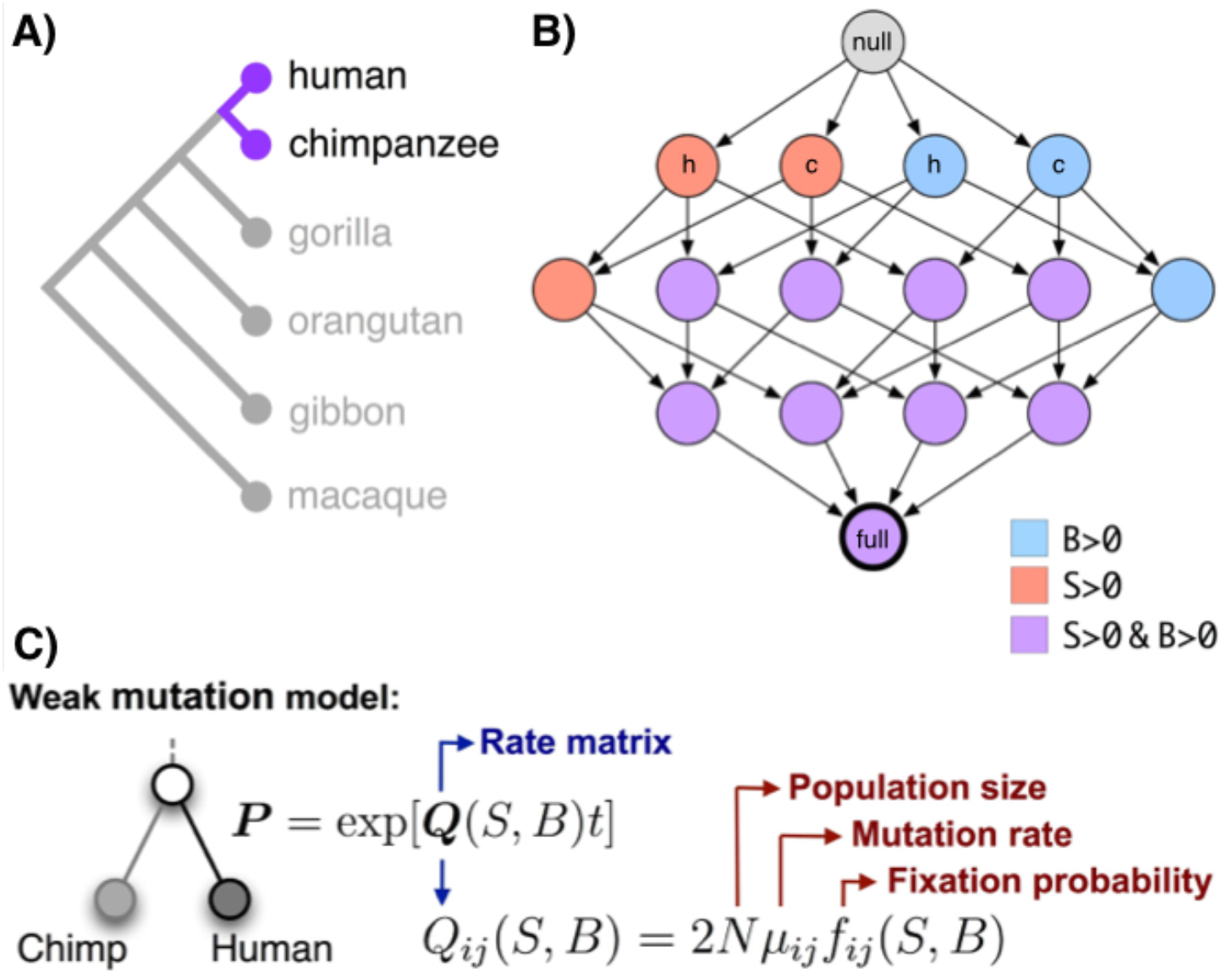
Testing for acceleration in five apes. To test a conserved non-coding element for an accelerated rate of unbiased and/or GC-biased substitutions in any of five apes, we perform a series of up to 1,024 nested likelihood ratio tests. **A)** The apes tested and their phylogenetic relationships (branches not to scale). For illustrating the approach, consider the possible models for just the human and chimpanzee branches. **B)** Example model selection procedure for two species. The null model (grey top node) has no unbiased acceleration (S=0 on both human and chimp branches) and no GC-biased acceleration (B=0 on both branches). There are four models with one of the parameters not equal to zero (next to top set of four nodes): S>0 in human (red, “h”), S>0 in chimp (red, “c”), B>0 in human (blue, “h”), and B>0 in chimp (blue, “c”). These are nested inside the six (i.e., 4 choose 2) possible models with two parameters constrained to zero, which are nested inside the four possible models with one parameter constrained to zero. The full model (purple, dark outlined bottom node) has all four parameters greater than zero. Our algorithm starts with the null model and performs the likelihood ratio test corresponding to each subsequent arrow moving from top to bottom of the graph of possible models, stopping if none of the models with an additional parameter greater than zero has significantly higher likelihood than the current model. For all five primates, the graph of nested models has 1,024 nodes and a similar structure. **C)** Each likelihood ratio test uses the weak mutation model of Kostka et al. (Kostka, Hubisz et al. 2012), illustrated here for one branch.

That is, to efficiently search for the best model we traverse the space of all alternative models in a specific way such that *ϑ_Aj_* has exactly one parameter more than *ϑ_Ai_*. In this case the asymptotic distribution of *Δ_ij_*(*X*) is a mixture of a *χ*^2^-distribution with one degree of freedom and a point mass of weight ½ at zero (Self and Liang 1987). This method will perform the LRT for a subset of models, built up from the best fitting models with acceleration in one lineage. Importantly, we stop traversing model space as soon as none of the more complex models considered yields a significantly better fit, compared with the best-fitting model nested within them. This produces a set of lineages for which we have evidence of a lineage-specific substitution rate increase. Finally, we annotate each alignment *X* with a single representative model: If all models we scored are nested within each other, we report the most complex model that fits the alignment better than all simpler models with fewer parameters (i.e., we use the last significantly better model from the forward selection). If the selection procedure generates some models that are not perfectly nested, we select the model with the best Akaike Information Criterion (AIC), as a trade off between model complexity and model fit.

This procedure outputs a single model per alignment block, which is either the null model or the best alternative model that represents a combination of acceleration (unbiased or GC-biased) across lineages that is significantly more likely than the null model given the alignment. Additional details are provided in the Methods and Supplementary Text.

### Implementation

The model selection method is implemented as an open-source software package in **R**, called linACC (code available at http://www.kostkalab.net/software.html). The package is based on methods implemented in the RPHAST package (Hubisz, Pollard et al. 2011) (http://compgen.cshl.edu/phast/).

## Results

### Apes have lineage-specific accelerated regions

We analyzed genome-wide multiple sequence alignments of 100 vertebrates and identified 272,466 mammalian conserved elements that met our stringent quality criteria (Methods). We excluded the apes we test for acceleration in defining these conserved elements. Each element was then evaluated for accelerated substitution rates in the lineages leading to human, chimpanzee, gorilla, orangutan, and gibbon from their most recent common ancestor with another ape (**Figure 1a**). Comparing substitution rates in apes to those expected given a null model of conservation similar to that seen in other apes allows us to detect unbiased acceleration (consistent with positive selection or loss of constraint) and GC-biased acceleration (consistent with gBGC) in any combination of the ape lineages. To statistically compare the partially nested set of 1,024 possible combinations of unbiased and/or GC-biased acceleration in any of the lineages, we designed and performed an efficient forward model selection procedure (see above and Methods). We identified 5,916 mammalian conserved genomic elements with statistically significant evidence of accelerated substitutions in at least one ape lineage (p<0.0001; LRT of the best model compared to the null model). We chose this somewhat conservative p-value threshold because formally controlling a multiple testing error rate in the context of our model selection procedure is not straightforward. These lineage-specific accelerated regions (linARs) have more substitutions than expected in various combinations of all five lineages. The human-accelerated linARs overlap previously identified HARs as much as expected based on prior comparisons of HARs derived with different data and methods (**Supplemental Text**) (Franchini and Pollard 2017). We describe relative numbers for different apes below.

### Genomic distribution of ape linARs

Similar to HARs, ape linARs are mostly non-coding with the majority falling in intergenic regions of the human genome (**Table 1**). This distribution is similar to that of the phastCons elements from which linARs are drawn, except that linARs are more enriched for intergenic elements (45.6% of linAR sequence versus 29.1% of phastCons sequence; p<0.001) (**Figure 2**). This trend is consistent across linARs that are accelerated in different apes, and it holds for both unbiased and GC-biased acceleration (**Supplemental Figure S1, Table 1**). Interestingly, linARs show clustering along human chromosomes, as was previously observed with HARs (**Supplemental Figure S2**). HARs are significantly enriched at the distal ends of human chromosomes, where substitution and recombination rates are elevated (Pollard, Salama et al. 2006, Kamm, Pisciottano et al. 2013). But this pattern appears weaker for all ape linARs in human genome coordinates, likely due in part to chromosomal rearrangements that moved ancestrally distal chromosomal segments into non-distal regions of the human chromosomes.

**Figure 2.**
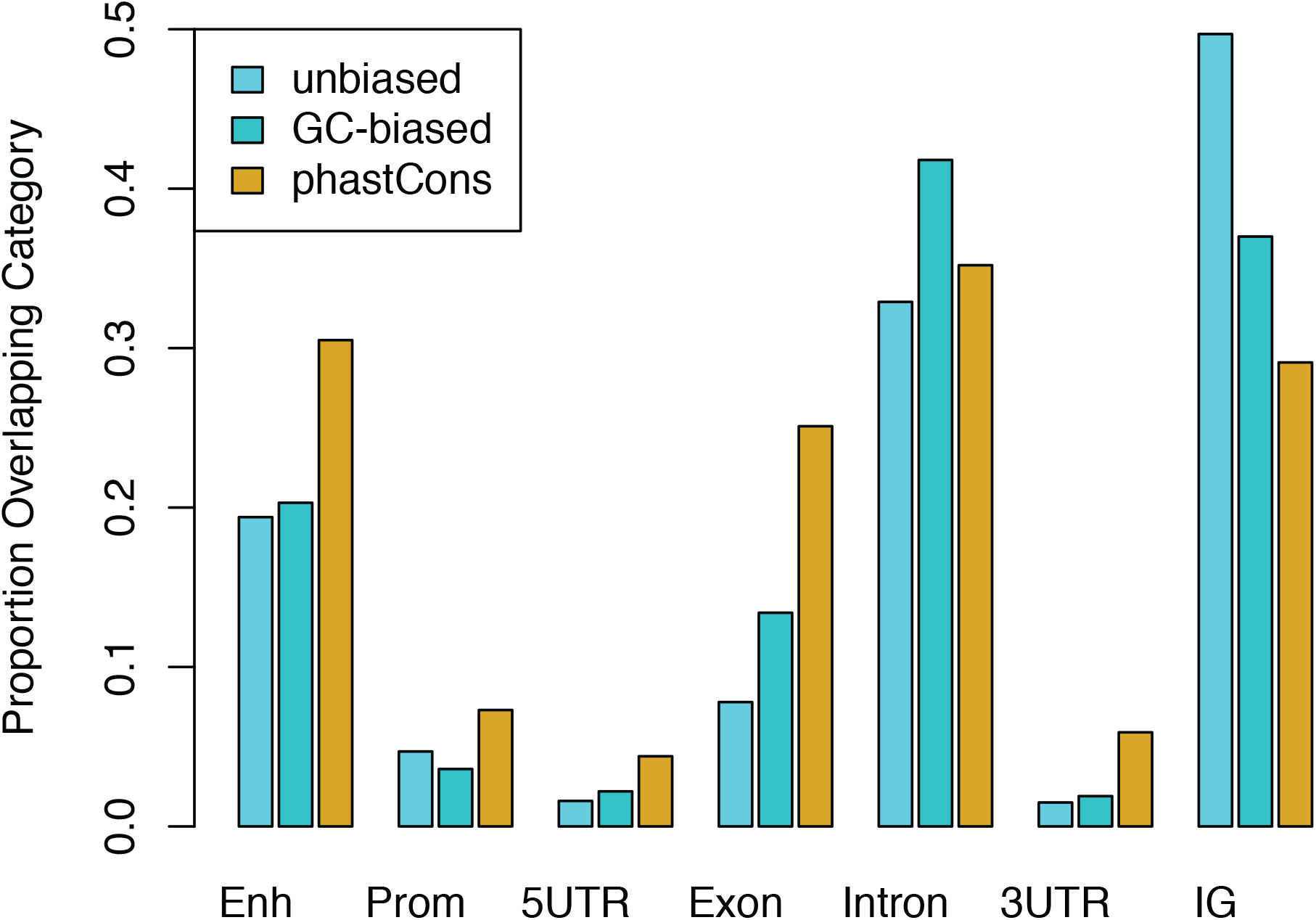
linARs are enriched in intergenic regions. For each genomic feature annotation category, bar height is the proportion of linARs (unbiased acceleration, light blue; GC-biased acceleration, turquoise) and all phastCons elements (yellow) overlapping that feature. Enh: enhancer, Prom: promoter, 5UTR: 5’ untranslated region, 3UTR: 3’ untranslated region, IG: intergenic.

**Table 1.** Proportions of the base pairs in linARs and all phastCons elements that overlap various genomic features, allowing for multiple overlapping annotations show enrichment of linARs in intergenic regions. Accel = acceleration type: unbiased acceleration is S>0; GC-biased acceleration is B>0.

| Type of linAR |  | Proportion of linAR base pairs in each category<br>(allowing multiple categories if overlapping) |  |  |  |  |  |  |
| --- | --- | --- | --- | --- | --- | --- | --- | --- |
| Species | Accel | Enhancer | Promoter | 5' UTR | Exon | Intron | 3' UTR | Intergenic |
| Human | S>0 | 0.219 | 0.027 | 0.004 | 0.045 | 0.361 | 0.016 | 0.482 |
| Chimp | S>0 | 0.215 | 0.066 | 0.032 | 0.139 | 0.329 | 0.049 | 0.422 |
| Gorilla | S>0 | 0.196 | 0.037 | 0.012 | 0.053 | 0.329 | 0.018 | 0.509 |
| Orangutan | S>0 | 0.235 | 0.057 | 0.018 | 0.108 | 0.317 | 0.017 | 0.438 |
| Gibbon | S>0 | 0.194 | 0.047 | 0.016 | 0.078 | 0.329 | 0.015 | 0.497 |
| Any | S>0 | 0.223 | 0.053 | 0.020 | 0.104 | 0.335 | 0.028 | 0.448 |
| Human | B>0 | 0.271 | 0.085 | 0.031 | 0.075 | 0.381 | 0.017 | 0.420 |
| Chimp | B>0 | 0.292 | 0.043 | 0.012 | 0.09 | 0.385 | 0.028 | 0.407 |
| Gorilla | B>0 | 0.284 | 0.045 | 0.005 | 0.077 | 0.367 | 0.037 | 0.460 |
| Orangutan | B>0 | 0.204 | 0.018 | 0.010 | 0.171 | 0.343 | 0.031 | 0.437 |
| Gibbon | B>0 | 0.203 | 0.036 | 0.022 | 0.134 | 0.418 | 0.019 | 0.370 |
| Any | B>0 | 0.254 | 0.05 | 0.020 | 0.101 | 0.381 | 0.021 | 0.420 |
| Any | S>0, B>0 | 0.220 | 0.046 | 0.017 | 0.088 | 0.348 | 0.023 | 0.456 |
| PhastCons | - | 0.305 | 0.073 | 0.044 | 0.251 | 0.352 | 0.059 | 0.291 |

### Ape linARs may function as gene regulatory enhancers

The genomic distribution and evolutionary conservation (outside apes) of non-coding linARs is suggestive of regulatory function. To explore this hypothesis, we annotated phastCons elements, including linARs, with a wide variety of publicly available data, including functional genomics (ChIP-seq and RNA-seq) data and the VISTA Enhancer Browser (**Supplemental Table S1**). We found strong support for a regulatory function for the majority of linARs, with 75.6% of linARs containing enhancer-associated marks (histone 3 lysine 27 acetylation, p300 binding) or enhancer predictions in human and/or mouse cells. We also found that 58/117 (49.6%) linARs tested by VISTA show evidence of enhancer activity in mouse embryos, which is ~1.26-fold more than expected given the VISTA validation rate of phastCons elements (binomial p=0.0166). Since much of this annotation is based on human sequences and/or human cells, further studies are needed to determine if the putative regulatory functions of linARs are conserved across apes.

### Developmental loci are enriched for ape linARs

Given the potential regulatory role of linARs, we were curious about the functions of genes regulated by linARs. We therefore mapped each linAR to the nearest gene and tested if genes associated with linARs are enriched for any gene ontology categories compared to phastCons elements. We found a strong enrichment for genes involved in developmental processes, in particular central nervous system development, as well as functions related to transcription factor activity (**Table 2, Supplemental Table S2**). This pattern is consistent across linARs from different apes (**Supplemental Tables S3–S7**) and similar to the functional enrichments previously reported for HARs (Lindblad-Toh, Garber et al. 2011), demonstrating a shared link between accelerated sequence evolution and developmental processes across apes. The enrichment of developmental regulators amongst linAR-associated genes is robust to the bioinformatics method for mapping phastCons elements to genes (Methods). The enrichment is weaker, however, when analyzed from the perspective of the phastCons elements (linAR vs. not) rather than the perspective of the genes (linAR-associated vs. not). This difference is driven in part by genes with large regulatory domains that harbor many phastCons elements. Thus, linARs frequently occur close together on the genome nearby developmental transcription factors, but these loci also harbor many non-accelerated conserved non-coding sequences.

**Table 2.** Gene Ontology (GO) enrichments for nearest genes to linARs compared to the background of phastCons elements, performed using GREAT (McLean, Bristor et al. 2010) to map linARs to nearest genes and GOrilla to compute enrichments. (Eden, Lipson et al. 2007, Eden, Navon et al. 2009). Only the top hits are shown; see Table S2 for a full list of enriched GO terms.

| **GO term** | **Description** | **FDR** | **Enrichment** |
| --- | --- | --- | --- |
| ***Biological Process*** |  |  |  |
| GO:2000026 | regulation of multicellular organismal development | 3.25E-7 | 1.28 |
| GO:0045595 | regulation of cell differentiation | 6.18E-7 | 1.29 |
| GO:0050793 | regulation of developmental process | 7.08E-7 | 1.24 |
| GO:0009653 | anatomical structure morphogenesis | 6.81E-6 | 1.29 |
| GO:0051960 | regulation of nervous system development | 6.41E-6 | 1.38 |
| GO:0007155 | cell adhesion | 5.64E-6 | 1.41 |
| GO:0022610 | biological adhesion | 8.94E-6 | 1.40 |
| ***Molecular Function*** |  |  |  |
| GO:0043565 | sequence-specific DNA binding | 2.06E-5 | 1.32 |
| GO:0000976 | transcription regulatory region sequence-specific DNA binding | 3.14E-4 | 1.35 |
| GO:1990837 | sequence-specific double-stranded DNA binding | 2.97E-4 | 1.34 |
| GO:0044212 | transcription regulatory region DNA binding | 5.67E-4 | 1.31 |
| GO:0001067 | regulatory region nucleic acid binding | 5.69E-4 | 1.30 |
| ***Cellular Component*** |  |  |  |
| GO:0030426 | growth cone | 5.03E-2 | 1.69 |
| GO:0097060 | synaptic membrane | 4.38E-2 | 1.45 |
| GO:0030427 | site of polarized growth | 8.07E-2 | 1.61 |

### Hotspots of accelerated evolution within and across ape species

Because linARs are clustered in the human genome (**Supplemental Figure S3**), we sought to identify specific genomic regions with large clusters of linARs. First, we considered each ape separately. For each species, we compared the median distance between closest linARs for that species to the same statistic computed on random sets of equal numbers of phastCons elements (Methods). This analysis showed that species-specific linARs are significantly more clustered than phastCons elements for each species alone: human (p<0.001), chimpanzee (p=0.002), orangutan (p<0.001), gorilla (p<0.001), and gibbon (p<0.001). Since phastCons elements are themselves fairly clustered, we conclude that species-specific linARs show strong clustering in all five apes.

We next sought to compare the genomic distribution of linARs across species. Using human genome coordinates, we repeated the statistical test for distance to the nearest linAR including all 5,916 linARs. This revealed that linARs are closer together on average than expected given the genome-wide distribution of phastCons elements (p<0.001) (**Figure 3**). To identify discrete linAR clusters, we first clustered phastCons elements into groups for which the longest distance between consecutive elements is less than 100 kb. We found 175 (out of 1164) such clusters that contain more linARs than expected (false discovery rate (FDR) < 0.05; Binomial test) (**Supplemental Table S8**). Most clusters contain linARs from multiple different apes. Furthermore, all clusters are located in syntenic regions of the other ape genomes with their clustering preserved (Methods). Thus, not only do ape linARs cluster within species, but these clusters are also maintained across species.

**Figure 3.**
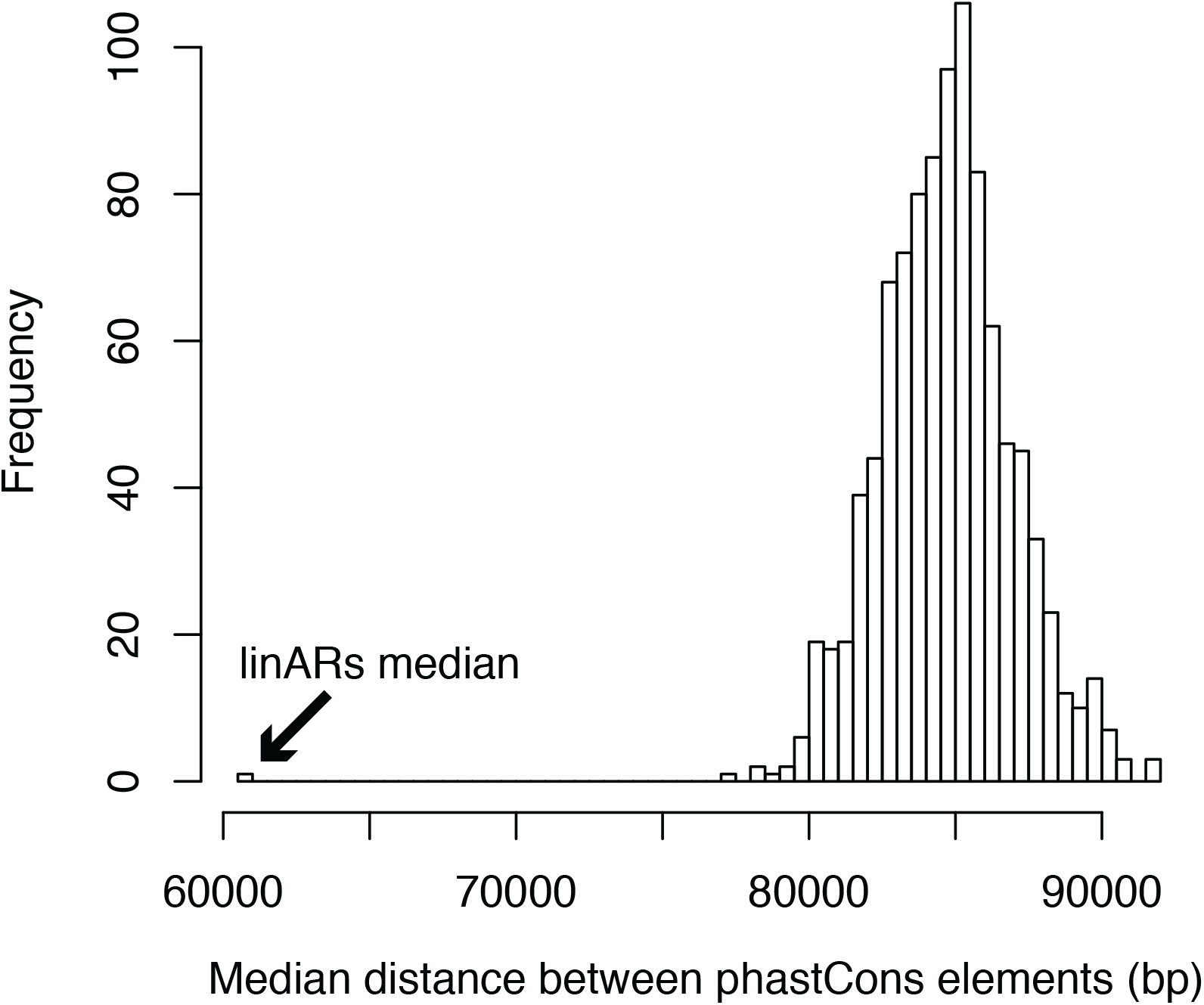
The linARs are more clustered than phastCons elements. The distribution of median distances between phastCons elements was computed using 1000 random draws of 5,916 phastCons elements from the full set. The median distance is >80 kb for most sets of 5,916 phastCons elements (median = 84.7 kb), whereas the median distance between pairs of linARs is just 60.8 kb (arrow). This analysis shows that linARs are significantly closer to each other in the human genome than are random sets of the same number of phastCons elements (p<0.0001).

To explore the set of genes nearby linAR clusters, we mapped linAR clusters to any gene within the cluster boundaries or the closest gene if the cluster is intergenic and then ranked the resulting 219 genes based on the size of their clusters (**Supplemental Table S8**). Many of the top genes are developmental transcription factors and signaling proteins, including many expressed during development of the central nervous system and sensory organs. For example, one hotspot for linARs is a group of four clusters comprising 42 linARs that are located nearby each other in the locus of *ROBO1* and *ROBO2*, transmembrane genes that function as receptors for SLIT family proteins in axon guidance and cell migration. We also identified multi-species linAR clusters in the FOXP1 and FOXP2 loci. The largest cluster is a nearly 1.5-mb region on human chromosome 4 with 62 linARs, including elements accelerated in each ape. A potential gene target for these linARs is the neurodevelopmental regulator *TENM3* (**Figure 4b**). Supporting the hypothesis that regulation of *TENM3* evolved rapidly in different ape lineages, another large cluster of mixed-species linARs is located nearby *TENM2* (**Figure 4a**). These teneurin transmembrane proteins are co-expressed in neurodevelopment and can form a heterodimer. Finally, our analysis found a cluster of 36 linARs from multiple apes nearby the *NPAS3* gene, which was previously shown to harbor a cluster of HARs, several of which are validated neurodevelopmental enhancers (Kamm, Pisciottano et al. 2013). Together our results suggest that a discrete set of developmental regulatory loci have been subject to accelerated evolution independently in multiple ape lineages, suggesting that positive selection and other evolutionary processes preferentially targeted particular genes and pathways that regulate embryonic development.

**Figure 4.**
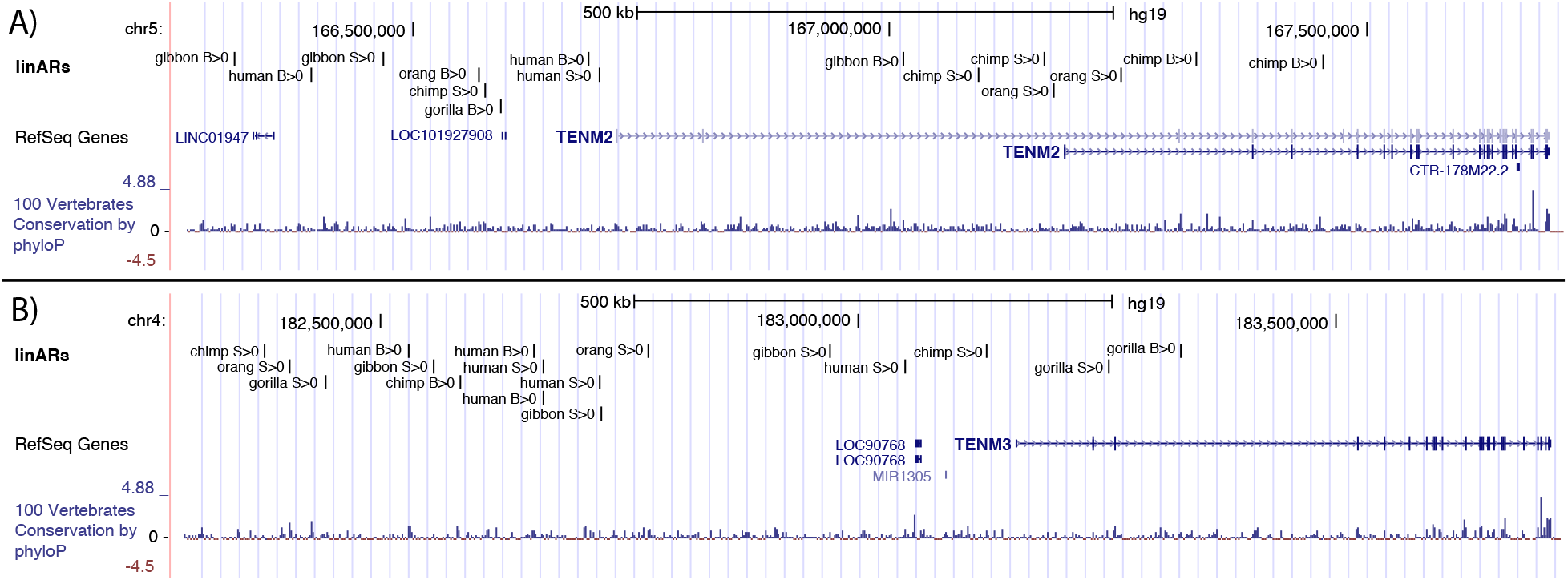
Clusters of linARs nearby teneurin transmembrane genes. Browser views of loci containing two of the largest multi-species linAR clusters nearby the genes **A)** *TENM2* and **B)** *TENM3*. Genes and conservation are shown with UCSC Genome Browser tracks. A custom track shows linARs annotated by species and unbiased acceleration (S>0) or GC-biased acceleration (B>0).

### Comparison of amounts of acceleration across apes

Our analyses in principle enable a direct comparison of the number of linARs across species. This comparison is confounded, however, by differences in the quality of genomes in the multiple sequence alignments we analyzed, including differences in sequence depth, assembly errors, and alignment artifacts, as well as the fact that the human assembly was used to scaffold some other genome assemblies. Supporting this confounding, the number of linARs with acceleration is negatively correlated with genome coverage, being lowest for human and gorilla. Accounting for this caveat, we find roughly equal numbers of linARs across the five ape lineages (**Figure 5**). This suggests that no ape genome, including the human genome, has a rate of lineage-specific acceleration that is qualitatively high compared to others.

**Figure 5.**
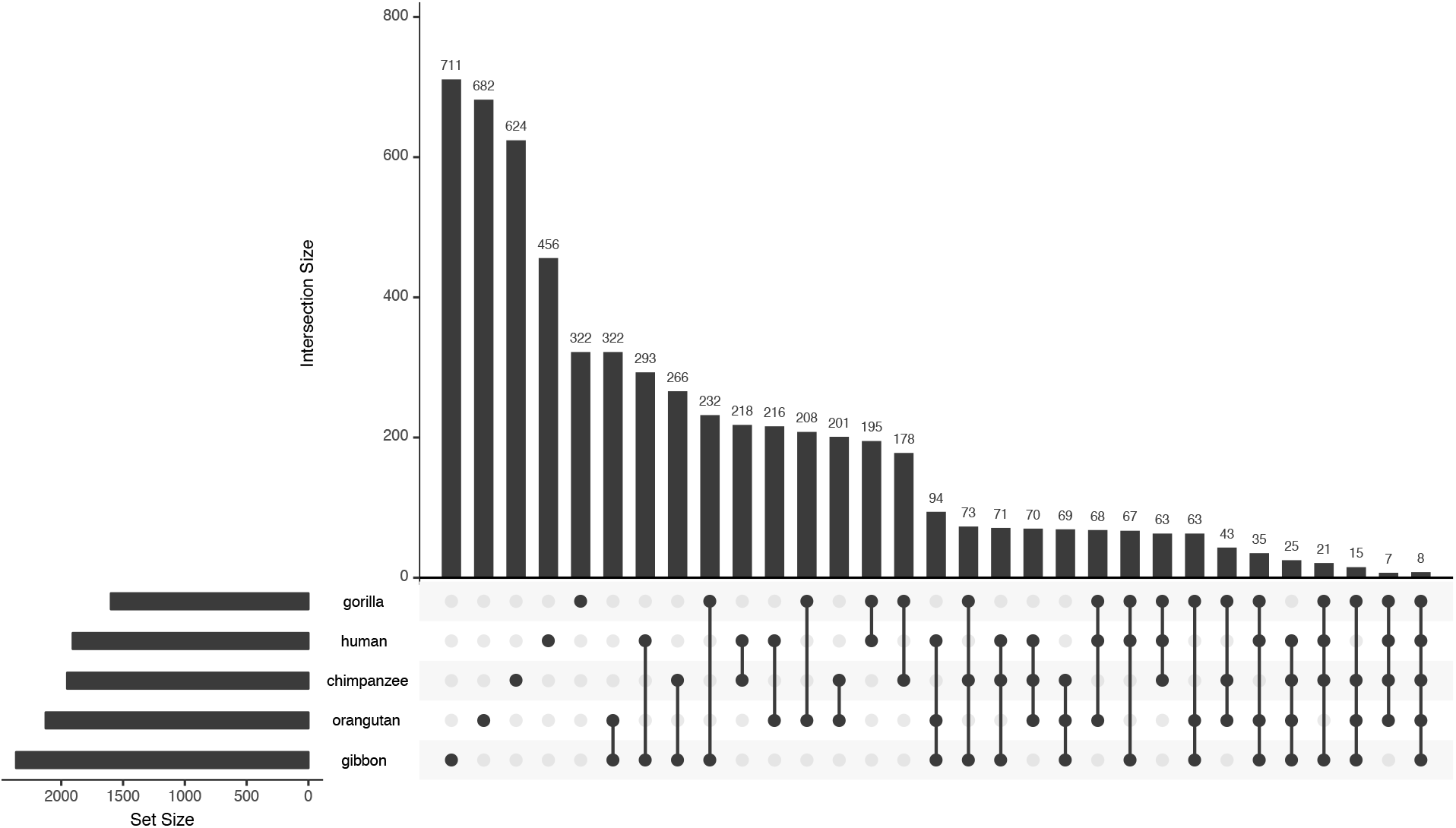
Apes have similar numbers of linARs. The upset plot (histogram) of linARs with acceleration on different subsets of lineages reveals three patterns: (i) most linARs are accelerated in one or few lineages, (ii) genomes with lower quality assemblies (e.g., gibbon) have the most linARs alone and in combination with acceleration in other lineages, and (iii) species otherwise have similar number of linARs along and in combination with each of the other apes.

Because we evaluated models with combinations of acceleration in multiple lineages, we could further evaluate patterns of co-occurrence of linARs on the ape phylogeny (**Figure 5**). Most linARs are accelerated in only one ape, with acceleration in two apes being next most common. Only 111 linARs have evidence of acceleration in four or five of the apes. Amongst the linARs with evidence of acceleration in two or three ape genomes, one of the species is commonly gibbon, which is consistent with faster evolution and/or lower genome quality for gibbon, both of which would increase the number of gibbon linARs and hence the probability of them overlapping linARs from other species. Besides cooccurrence with gibbon acceleration, linARs do not appear to be preferentially shared between sister species or other specific combinations of species. These results suggest that the evolutionary forces that accelerated the evolution of linARs typically affected just one lineage but also occasionally acted recurrently in the same genomic region multiple times during ape evolution, as has been observed in analyses of great ape population genetic diversity (Cagan, Theunert et al. 2016) and incomplete lineage sorting (Munch, Nam et al. 2016).

### Evidence of biased gene conversion

Our statistical models include separate parameters for unbiased acceleration (consistent with positive selection or loss of constraint) and GC-biased acceleration (consistent with gBGC or selection on GC-content). In each lineage, the model selection procedure directly compares the likelihood of models with either or both of these parameters. This enables us to quantify the relative rates of unbiased and GC-biased acceleration across species. All apes had many more linARs with unbiased acceleration, although GC-biased acceleration was not uncommon (26.2% of linARs overall, range 17.7-32.3% per ape) (**Table 3**), consistent with estimates of the prevalence of unbiased positive selection in previous studies of human accelerated non-coding regions (Kostka, Hubisz et al. 2012, Gittelman, Hun et al. 2015). This suggests that gBGC is prevalent in apes, albeit with differences in frequency, but it is consistently less common than the combination of positive selection and loss of constraint.

**Table 3.** Counts of linARs with unbiased and/or GC-biased acceleration in different combinations of ape lineages. Chimp = chimpanzee, Orang = orangutan, S>0 = unbiased acceleration, B>0 = GC-biased acceleration. Only pairs of lineages are shown; a small number of linARs are accelerated in more than two lineages.

|  | Human<br>S>0 | Human<br>B>0 | Chimp<br>S>0 | Chimp<br>B>0 | Gorilla<br>S>0 | Gorilla<br>B>0 | Orang<br>S>0 | Orang<br>B>0 | Gibbon<br>S>0 | Gibbon<br>B>0 |
| --- | --- | --- | --- | --- | --- | --- | --- | --- | --- | --- |
| Human<br>S>0 | 1,298 | 9 | 257 | 80 | 260 | 70 | 310 | 61 | 343 | 83 |
| Human<br>B>0 |  | 618 | 95 | 53 | 80 | 55 | 114 | 39 | 122 | 67 |
| Chimp<br>S>0 |  |  | 1,426 | 12 | 197 | 64 | 250 | 62 | 294 | 88 |
| Chimp<br>B>0 |  |  |  | 538 | 89 | 58 | 103 | 28 | 127 | 44 |
| Gorilla<br>S>0 |  |  |  |  | 1,091 | 3 | 273 | 49 | 285 | 71 |
| Gorilla<br>B>0 |  |  |  |  |  | 510 | 97 | 31 | 114 | 47 |
| Orang<br>S>0 |  |  |  |  |  |  | 1,764 | 18 | 412 | 110 |
| Orang<br>B>0 |  |  |  |  |  |  |  | 380 | 83 | 34 |
| Gibbon<br>S>0 |  |  |  |  |  |  |  |  | 1,814 | 24 |
| Gibbon<br>B>0 |  |  |  |  |  |  |  |  |  | 575 |

## Discussion

We developed a model selection procedure to scan ape genomes in parallel for conserved elements similar to HARs. These analyses revealed that the human genome is not unique in having linARs, nor in how many linARs it has. Comparing patterns across five apes, we found fairly consistent numbers of linARs. Differences in counts of linARs (range 1,601-2,389) may be due primarily to variation in genome assembly quality. Another bias to consider in interpreting these results is the fact that the human genome was used as the reference genome in our analyses and was also employed in the assembly of other ape genomes. Across species, we also found fairly similar proportions of unbiased versus GC-biased linARs (mean 26.2%). Variation in the GC-biased proportion (range 17.7-32.2%) could potentially reflect differences in rates and patterns of gBGC versus positive selection or loss of constraint between apes.

Another striking similarity of linARs across apes is that they cluster together (64% within 100 kb of another linAR), both within and between species, in loci harboring developmental genes (enrichment=1.1, FDR<0.0004). These are mostly distinct non-coding elements in the same locus, though some linARs are accelerated in two or more of the ape lineages. An intriguing question for future research is to determine the mechanisms driving the clustering of linARs. Here we analyzed linARs in comparison to phastCons clusters in the human genome (reference sequence in alignments) and eliminated the possibility that linARs clustering is simply due to the higher number of conserved elements in developmental loci or the larger intergenic distances in these loci, which results in more conserved elements being closest to a developmental gene. While the full set of conserved elements did show clustering, linARs were more densely clustered than other conserved elements and more frequently associated with developmental loci. One possibility is that recurrent selection on the expression levels of certain developmental genes has occurred throughout primate evolution, which is consistent with other studies that found evidence of recurrent selection in the ape species we analyzed (Cagan, Theunert et al. 2016, Munch, Nam et al. 2016). Alternatively, certain genes may tolerate more regulatory evolution (though developmental gene expression tends to be deeply conserved (Li, Huang et al. 2014)). Another hypothesis is that the genomic regions containing linARs are mutational hotspots and that loss of constraint, rather than positive selection, drives the clusters. Previous work found no evidence in support of elevated mutation rates in 40 kb regions nearby 49 HARs (Katzman, Kern et al. 2010), but future studies are needed to confirm if this is true for linARs more broadly.

Ape linARs are particularly enriched nearby developmental transcription factors and other regulators of embryonic development, as was previously observed for HARs (Lindblad-Toh, Garber et al. 2011). This suggests the tantalizing hypothesis that mutations in linARs altered morphogenesis during ape evolution by modifying expression levels of key regulators of embryonic development. In future work, it would be interesting to test if lineage-specific changes in the sequences of non-human ape linARs alter gene expression during embryonic development, as has been demonstrated for several HARs (Hubisz and Pollard 2014, Boyd, Skove et al. 2015). If the linARs that cluster nearby a developmental regulator are enhancers that control distinct spatial or temporal aspects of that gene’s expression, then one could hypothesize that their accelerated evolution in different apes might be associated with morphological features that diverged during ape evolution. On the other hand, the multiple regulatory elements nearby a developmental gene may also buffer or otherwise affect each other (Long, Prescott et al. 2016), making it hard to predict the effect that sequence changes in one small non-coding element will have on gene expression and organismal phenotypes. These questions will be best answered using functional studies that test linARs not only individually, but also collectively. These investigations may soon be possible with high-throughput technologies, such as genome editing and massively parallel reporter assays, which enable thousands of regulatory elements to be investigated en masse in primate cells. With these approaches, the role of linARs in primate evolution and their clustering in specific developmental pathways could soon be elucidated.

## Material and Methods

Additional details about methods are available in the supplemental text, our open-source R-package linACC that implements model selection (available at *www.kostkalab.net/software.html*). Our methods and open source software extend the methods in RPHAST (Hubisz, Pollard et al. 2011). We also posted most of our analysis scripts plus accompanying data sets online at: *http://www.kostkalab.net/pubsoftware/linACC/supplement/linACCsupplement.html*.

### Sequence data

We obtained **multiz** whole-genome sequence alignments of 100 vertebrates and associated phylogenetic trees for the autosomes and chromosome X from the UCSC Genome Browser (http://hgdownload.soe.ucsc.edu/goldenPath/hg19/multiz100way/). The reference genome for the alignments is human (hg19 assembly).

### Conservation

To analyze a consistent set of genomic elements that are likely functional, we used the **phastCons** program to identify mammalian conserved elements genome-wide while excluding all apes from the computations (human reference sequence masked and other apes dropped from the alignments). Command line parameters were: --rho 0.3 --expected-length 45 --target-coverage 0.3. These are the standard parameters used in the UCSC Genome Browser protocol. The genome was analyzed in 10 megabase (Mb) blocks to facilitate computations. Conserved elements separated by less than 10 base pairs (bp) were merged, and then any elements shorter than 50 bp were dropped, since power to detect ape acceleration is low on short alignments (Pollard, Hubisz et al. 2010). We also dropped any element where any of the five apes (human, chimpanzee, gorilla, orangutan, and gibbon) was not present in the alignment (i.e., at least one site with a nucleotide). This produces 660,077 conserved elements that cover 94,370,847 bp of the human genome. Alignments corresponding to each conserved element were extracted from the 100-species genome-wide alignments.

### Filtering

To minimize the influence of sequencing, assembly, and alignment errors on our inferences regarding accelerated substitution rates, we filtered the conserved element alignments using an extension of a previous approach (Lindblad-Toh, Garber et al. 2011). We first masked repeat sequences (UCSC srpt and rmsk tracks) annotated in each of the five ape genomes that we tested for accelerated evolution (hg19, panTro4, gorGor3, ponAbe2, nomLeu3). Next, we generated genome-wide self-alignments following the UCSC selfChain documentation: align to self using **lastz**, and then chain into longer contiguous alignments. Bases in these self-similar regions were masked in the conserved element alignments. We then dropped alignments corresponding to annotated pseudogenes (pseudoYale60 UCSC Genome Browser track), segmental duplications (genomicSuperDups track), and self-similar genomic regions (selfChain, see above) annotated on the reference genome (hg19). Finally, we used the UCSC netSyntenic tracks to require that conserved elements fall within blocks of level-1 or level-2 non-gapped synteny between human (hg19) and (i) macaque (rheMac3), (ii) dog (canFam3), and (iii) mouse (mm10). Together, these conservative filters likely remove some truly accelerated elements, but they are necessary to avoid thousands of false inferences of accelerated evolution due to misaligned paralogous sequences and other errors that are present in genome-wide alignments (Pollard, Salama et al. 2006, Lindblad-Toh, Garber et al. 2011). This bioinformatics pipeline generated multiple sequence alignments for 272,466 conserved elements (“phastCons elements”) covering 37,152,199 bp of the human genome that are our candidates for accelerated evolution in apes.

### Testing for acceleration in ape lineages

We applied our model selection procedure (see above) to multiple sequence alignments of the 272,466 high-quality phastCons elements to test for accelerated evolution in five apes: human, chimpanzee, gorilla, orangutan, and gibbon. For five lineages, there are 2^2×5^=1,024 different alternative models *ϑ_Ai_* per alignment, which we efficiently screened with our forward selection algorithm.

At the first step of forward selection, all models {*ϑ_Ai_*} with one accelerated lineage (either unbiased or GC-biased) are compared to the null model with no acceleration (*ϑ_0_*) using a likelihood ratio test (LRT) on the filtered multiple sequence alignment for a given phastCons element, and all models that fit significantly better than the null model (i.e., the ones with p-value less than 0.01) are retained. At each subsequent step in the forward selection, models with acceleration on additional lineages are compared to the retained simpler, nested models using an LRT. At each step significant models move forward to the next step of forward selection by adding all possible single additional lineages with acceleration (unbiased or GC-biased). The algorithm stops when adding more accelerated lineages does not provide any significant improvement in fit. After a final model selection step (see Results) it outputs a single best model per phastCons element alignment as well as the p-value from the LRT comparing the best model to the null model with no acceleration.

### Annotation

We explored the genomic distribution and potential functions of the resulting ape linARs using the UCSC known genes annotation (http://genome.ucsc.edu), the VISTA Enhancer Browser (http://enhancer.lbl.gov), ChromHMM genome segmentations of ENCODE data (https://www.encodeproject.org), FANTOM5 enhancers (http://fantom.gsc.riken.jp/5/), and other functional genomics data from publicly available databases (**Supplemental Table S1**). Each phastCons element (linARs and non-linARs) was annotated with all overlapping data, and annotation patterns were compared between linARs and all phastCons elements.

### Ontology

To test if linARs are preferentially associated with particular genes, we mapped each phastCons element (linARs and non-linARs) to the closest gene. These closest genes were used to compare gene ontology categories for genes associated with linARs versus genes associated with a phastCons element using GOrilla (Eden, Lipson et al. 2007, Eden, Navon et al. 2009) (http://cbl-gorilla.cs.technion.ac.il) with default settings. Genes associated with phastCons elements were identified using a random subset of 20,000 elements for computational efficiency. We also used GREAT (McLean, Bristor et al. 2010) (http://bejerano.stanford.edu/great/public/html/) to perform enrichment analyses that are non-coding element based, rather than gene based, and therefore account for the unequal size of regulatory domains of genes. To assess robustness, enrichment analyses were repeated using other methods of mapping phastCons elements to genes: two nearest genes and the basal-plus-extension method implemented in GREAT.

### Clustering

We investigated whether linARs within and across apes occur closer together along the human genome than expected given the density of phastCons elements. For each linAR, we computed the genomic distance to the nearest other linAR. Then we did the same for all phastCons elements. To test if the median distance between linARs is shorter than expected given the distances between phastCons elements, we randomly sampled (without replacement) 1,000 sets of phastCons elements of the same size as the number of linARs and computed the median distance between phastCons elements for each set. The proportion of these median distances that exceeded the median distance between linARs is the empirical p-value.

As a second approach, we identified clusters of phastCons elements where consecutive elements are separated by no more than 100 kilobases (kb) in the human genome (hg19, reference sequence in alignments). For each cluster containing at least one linAR, we calculated the number *n* of phastCons elements and the number *k* of those that are linARs. We then computed a Binomial p-value Bin(*k*|*p, n*) for the cluster containing *k* linARs out of *n* phastCons elements, under the null hypothesis that linARs are not clustered any more than typical phastCons elements (*i.e*., the probability *p* of a phastCons element in the cluster being a linAR was set equal to the overall proportion of linARs among phastCons elements genome-wide). We accounted for the fact that each cluster has at least one linAR by dividing the resulting p-values by 1-Bin(0 |*p,n*) before performing multiple testing correction to control the FDR (Hochberg and Benjamini 1990). This enrichment test yielded a set of clusters with more linARs than expected.

To evaluate synteny of linAR clusters in the non-human ape genomes, we used UCSC level 1 and 2 syntenic net tables (hg19 assembly).

## Acknowledgements

This work was supported by NIGMS (grant #GM82901), the San Simeon Fund, and the Gladstone Institutes. DK was also supported by NIGHM (grant #GM115836) and the University of Pittsburgh School of Medicine.

## References

Bird, C. P., B. E. Stranger, M. Liu, D. J. Thomas, C. E. Ingle, C. Beazley, W. Miller, M. E. Hurles and E. T. Dermitzakis (2007). “Fast-evolving noncoding sequences in the human genome.” Genome Biol 8(6): R118.

Booker, B. M., T. Friedrich, M. K. Mason, J. E. VanderMeer, J. Zhao, W. L. Eckalbar, M. Logan, N. Illing, K. S. Pollard and N. Ahituv (2016). “Bat Accelerated Regions Identify a Bat Forelimb Specific Enhancer in the HoxD Locus.” PLoS Genet 12(3): e1005738.

Boyd, J. L., S. L. Skove, J. P. Rouanet, L. J. Pilaz, T. Bepler, R. Gordan, G. A. Wray and D. L. Silver (2015). “Human-chimpanzee differences in a FZD8 enhancer alter cell-cycle dynamics in the developing neocortex.” Curr Biol 25(6): 772–779.

Bush, E. C. and B. T. Lahn (2008). “A genome-wide screen for noncoding elements important in primate evolution.” BMC Evol Biol 8: 17.

Cagan, A., C. Theunert, H. Laayouni, G. Santpere, M. Pybus, F. Casals, K. Prufer, A. Navarro, T. Marques-Bonet, J. Bertranpetit and A. M. Andres (2016). “Natural Selection in the Great Apes.” Mol Biol Evol 33(12): 3268–3283.

Capra, J. A., G. D. Erwin, G. McKinsey, J. L. Rubenstein and K. S. Pollard (2013). “Many human accelerated regions are developmental enhancers.” Philos Trans R Soc Lond B Biol Sci 368(1632): 20130025.

Eden, E., D. Lipson, S. Yogev and Z. Yakhini (2007). “Discovering motifs in ranked lists of DNA sequences.” PLoS Comput Biol 3(3): e39.

Eden, E., R. Navon, I. Steinfeld, D. Lipson and Z. Yakhini (2009). “GOrilla: a tool for discovery and visualization of enriched GO terms in ranked gene lists.” BMC Bioinformatics 10: 48.

Franchini, L. F. and K. S. Pollard (2017). “Human evolution: the non-coding revolution.” BMC Biol 15(1): 89.

Galtier, N. and L. Duret (2007). “Adaptation or biased gene conversion? Extending the null hypothesis of molecular evolution.” Trends Genet 23(6): 273–277.

Gittelman, R. M., E. Hun, F. Ay, J. Madeoy, L. Pennacchio, W. S. Noble, R. D. Hawkins and J. M. Akey (2015). “Comprehensive identification and analysis of human accelerated regulatory DNA.” Genome Res 25(9): 1245–1255.

Hochberg, Y. and Y. Benjamini (1990). “More powerful procedures for multiple significance testing.” Stat Med 9(7): 811–818.

Holloway, A. K., D. J. Begun, A. Siepel and K. S. Pollard (2008). “Accelerated sequence divergence of conserved genomic elements in Drosophila melanogaster.” Genome Res 18(10): 1592–1601.

Holloway, A. K., B. G. Bruneau, T. Sukonnik, J. L. Rubenstein and K. S. Pollard (2016). “Accelerated Evolution of Enhancer Hotspots in the Mammal Ancestor.” Mol Biol Evol 33(4): 1008–1018.

Hubisz, M. J. and K. S. Pollard (2014). “Exploring the genesis and functions of Human Accelerated Regions sheds light on their role in human evolution.” Curr Opin Genet Dev 29: 15–21.

Hubisz, M. J., K. S. Pollard and A. Siepel (2011). “PHAST and RPHAST: phylogenetic analysis with space/time models.” Brief Bioinform 12(1): 41–51.

Kamm, G. B., F. Pisciottano, R. Kliger and L. F. Franchini (2013). “The developmental brain gene NPAS3 contains the largest number of accelerated regulatory sequences in the human genome.” Mol Biol Evol 30(5): 1088–1102.

Katzman, S., A. D. Kern, K. S. Pollard, S. R. Salama and D. Haussler (2010). “GC-biased evolution near human accelerated regions.” PLoS Genet 6(5): e1000960.

Kosiol, C., T. Vinar, R. R. da Fonseca, M. J. Hubisz, C. D. Bustamante, R. Nielsen and A. Siepel (2008). “Patterns of positive selection in six Mammalian genomes.” PLoS Genet 4(8): e1000144.

Kostka, D., M. J. Hubisz, A. Siepel and K. S. Pollard (2012). “The role of GC-biased gene conversion in shaping the fastest evolving regions of the human genome.” Mol Biol Evol 29(3): 1047–1057.

Li, J. J., H. Huang, P. J. Bickel and S. E. Brenner (2014). “Comparison of D. melanogaster and C. elegans developmental stages, tissues, and cells by modENCODE RNA-seq data.” Genome Res 24(7): 1086–1101.

Lindblad-Toh, K., M. Garber, O. Zuk, M. F. Lin, B. J. Parker, S. Washietl, P. Kheradpour, J. Ernst, G. Jordan, E. Mauceli, L. D. Ward, C. B. Lowe, A. K. Holloway, M. Clamp, S. Gnerre, J. Alfoldi, K. Beal, J. Chang, H. Clawson, J. Cuff, F. Di Palma, S. Fitzgerald, P. Flicek, M. Guttman, M. J. Hubisz, D. B. Jaffe, I. Jungreis, W. J. Kent, D. Kostka, M. Lara, A. L. Martins, T. Massingham, I. Moltke, B. J. Raney, M. D. Rasmussen, J. Robinson, A. Stark, A. J. Vilella, J. Wen, X. Xie, M. C. Zody, P. Broad Institute Sequencing, T. Whole Genome Assembly, J. Baldwin, T. Bloom, C. W. Chin, D. Heiman, R. Nicol, C. Nusbaum, S. Young, J. Wilkinson, K. C. Worley, C. L. Kovar, D. M. Muzny, R. A. Gibbs, T. Baylor College of Medicine Human Genome Sequencing Center Sequencing, A. Cree, H. H. Dihn, G. Fowler, S. Jhangiani, V. Joshi, S. Lee, L. R. Lewis, L. V. Nazareth, G. Okwuonu, J. Santibanez, W. C. Warren, E. R. Mardis, G. M. Weinstock, R. K. Wilson, U. Genome Institute at Washington, K. Delehaunty, D. Dooling, C. Fronik, L. Fulton, B. Fulton, T. Graves, P. Minx, E. Sodergren, E. Birney, E. H. Margulies, J. Herrero, E. D. Green, D. Haussler, A. Siepel, N. Goldman, K. S. Pollard, J. S. Pedersen, E. S. Lander and M. Kellis (2011). “A high-resolution map of human evolutionary constraint using 29 mammals.” Nature 478(7370): 476–482.

Long, H. K., S. L. Prescott and J. Wysocka (2016). “Ever-Changing Landscapes: Transcriptional Enhancers in Development and Evolution.” Cell 167(5): 1170–1187.

McLean, C. Y., D. Bristor, M. Hiller, S. L. Clarke, B. T. Schaar, C. B. Lowe, A. M. Wenger and G. Bejerano (2010). “GREAT improves functional interpretation of cis-regulatory regions.” Nat Biotechnol 28(5): 495–501.

Munch, K., K. Nam, M. H. Schierup and T. Mailund (2016). “Selective Sweeps across Twenty Millions Years of Primate Evolution.” Mol Biol Evol 33(12): 3065–3074.

Pollard, K. S., M. J. Hubisz, K. R. Rosenbloom and A. Siepel (2010). “Detection of nonneutral substitution rates on mammalian phylogenies.” Genome Res 20(1): 110–121.

Pollard, K. S., S. R. Salama, B. King, A. D. Kern, T. Dreszer, S. Katzman, A. Siepel, J. S. Pedersen, G. Bejerano, R. Baertsch, K. R. Rosenbloom, J. Kent and D. Haussler (2006).“Forces shaping the fastest evolving regions in the human genome.” PLoS Genet 2(10): e168.

Pollard, K. S., S. R. Salama, N. Lambert, M. A. Lambot, S. Coppens, J. S. Pedersen, S. Katzman, B. King, C. Onodera, A. Siepel, A. D. Kern, C. Dehay, H. Igel, M. Ares, Jr., P. Vanderhaeghen and D. Haussler (2006). “An RNA gene expressed during cortical development evolved rapidly in humans.” Nature 443(7108): 167–172.

Prabhakar, S., J. P. Noonan, S. Paabo and E. M. Rubin (2006). “Accelerated evolution of conserved noncoding sequences in humans.” Science 314(5800): 786.

Self, S. G. and K.-Y. Liang (1987). “Asymptotic Properties of Maximum Likelihood Estimators and Likelihood Ratio Tests under Nonstandard Conditions.” Journal of the American Statistical Association 82(398): 605–610.

